# Parallel Group ICA + ICA: Joint Estimation of Linked Functional Network Variability and Structural Covariation with Application to Schizophrenia

**DOI:** 10.1101/595017

**Authors:** Shile Qi, Jing Sui, Jiayu Chen, Jingyu Liu, Rongtao Jiang, Rogers Silva, Armin Iraji, Eswar Damaraju, Mustafa Salman, Dongdong Lin, Zening Fu, Dongmei Zhi, Jessica A. Turner, Juan Bustillo, Judith M. Ford, Daniel H. Mathalon, James Voyvodic, Sarah McEwen, Adrian Preda, Aysenil Belger, Steven G. Potkin, Bryon A. Mueller, Tulay Adali, Vince D. Calhoun

## Abstract

There is growing evidence that rather than using a single brain imaging modality to study its association with physiological or symptomatic features, the field is paying more attention to fusion of multimodal information. However, most current multimodal fusion approaches that incorporate functional magnetic resonance imaging (fMRI) are restricted to second-level 3D features, rather than the original 4D fMRI data. This trade-off is that the valuable temporal information is not utilized during the fusion step. Here we are motivated to propose a novel approach called “parallel group ICA+ICA” that incorporates temporal fMRI information from GICA into a parallel ICA framework, aiming to enable direct fusion of first-level fMRI features with other modalities (*e.g.* structural MRI), which thus can detect linked functional network variability and structural covariations. Simulation results show that the proposed method yields accurate inter-modality linkage detection regardless of whether it is strong or weak. When applied to real data, we identified one pair of significantly associated fMRI-sMRI components that show group difference between schizophrenia and controls in both modalities. Finally, multiple cognitive domain scores can be predicted by the features identified in the linked component pair by our proposed method. We also show these multimodal brain features can predict multiple cognitive scores in an independent cohort. Overall, results demonstrate the ability of parallel GICA+ICA to estimate joint information from 4D and 3D data without discarding much of the available information up front, and the potential for using this approach to identify imaging biomarkers to study brain disorders.

## 1. INTRODUCTION

Magnetic resonance imaging (MRI) has provided remarkable new insights into the structure and function of human brain. Acquisition of multimodal MRI from the same subject has been widely adopted in brain imaging researches, as different modalities represent different perspectives of the brain functional, structural or anatomical properties. Moreover, there is growing evidence suggesting that instead of using a single brain imaging modality to study the association with physiologic or pathological properties, researchers are paying more attention to fusion of multimodal information, a method that can take advantage of multiple imaging techniques, and to uncover the latent relationships that might be missed from single modality imaging analysis (Calhoun and Sui, 2016; Suk, et al., 2014; Zhang, et al., 2011). For instance, multimodal fusion can tell us how brain structure and function are impacted by psychopathology, and which structural or functional aspects of pathology could drive human behavior or cognition (Sui, et al., 2014).

However, most current multimodal fusion approaches (including joint independent component analysis (jICA) (Calhoun, et al., 2006), multi-set canonical correlation analysis (mCCA) (Correa, et al., 2010), parallel ICA (pICA) (Liu, et al., 2009), parallel ICA with reference (pICAR) (Chen, et al., 2013), mCCA+jICA, mCCAR+jICA (Qi, et al., 2018a), and linked ICA (Groves, et al., 2011), etc.) are restricted to second-level 3D fMRI features, e.g., fractional amplitude of low frequency fluctuations (fALFF) or regional homogeneity (ReHo) for fMRI (subject × imaging feature variables), rather than first-level 4D imaging features (subject × voxel × time points). The main reason for using second-level features in multimodal fusion is to provide a simpler subspace in which to link the multimodal data (Calhoun and Adali, 2009). While this provides a powerful framework for capturing multimodal information, the trade-off is that some essential information may be lost. For example, in fMRI related multimodal fusion analysis, the temporal dynamic information was not included in the above fusion methods.

It is well known that the temporal variation in fMRI signal conveys important information (Schmithorst and Holland, 2004). Reducing the 4D fMRI data to 3D spatial maps (non-temporal features) prior to the data fusion step does not allow the fusion process to utilize the temporal information. On the other hand, group ICA (GICA) (Calhoun, et al., 2001) is an approach that operates on first-level 4D fMRI data for multiple subjects, which is able to extract both group and subject-specific independent components (ICs) as well as their time courses. Although there are other ICA related method that can deal with fMRI, such as probabilistic ICA (Beckmann and Smith, 2004) and noisy ICA (Cichocki, et al., 1998; Griffanti, et al., 2017), GICA has demonstrated great potential to deal with multi-subject fMRI data, so in this paper our proposed method is based on GICA. GICA allows us to establish a correspondence of group ICs with subjects’ ICs while fully leveraging the temporal information. In contrast, parallel ICA aims to simultaneously identify ICs of two modalities and the linkage between them by maximizing both inter-modality correlation and intra-modality independence. Building on the success of GICA to leverage temporal information as well as the flexible fusion framework of parallel ICA, we combine GICA and pICA in order to simultaneously analyze both first-level fMRI and structural MRI (sMRI) features, with the purpose of identifying linked functional spatial network variability and structural covariations, while retaining the original desirable properties of both pICA and GICA. Therefore, we propose to extend the pICA method and link brain structure to first-level functional MRI data via direct optimization of their associations, enabling us to reveal structural influence on coherent functional network variability.

Parallel GICA+ICA works by defining a variability matrix that measures the subject-level variations between group and back-reconstructed subject spatial maps within GICA, and maximizing the correlation of that measure with subject expression profiles from an ICA decomposition of a sMRI dataset. In order to achieve data fusion between structural (subject × voxel-wise gray matter (GM) volume values) and functional data (subject × voxel × time points), a subject-level summary feature that captures variability among subjects must be defined for fMRI, so that direct associations can be measured between both modalities. To that end, we note that the group-level ICs from GICA capture spatial co-activation patterns shared among all the subjects, each forming an independent brain network. In this sense, group ICs can be interpreted as a cohort common pattern representing a functional brain network template shared over all subjects. Therefore, it is worthwhile to examine how much the subject-specific ICs deviate from the shared common pattern and whether this deviation may serve as a summarized fMRI feature which associates with structural data (Chen, et al., 2018). Based on this feature, a novel parallel GICA+ICA approach that leverages the first-level temporal information from fMRI can be derived enabling the discovery of associations in the form of linked covariation between functional spatial network patterns and structural features. The key difference from pICA is that an intermediate subject-level variability matrix (***C***_1_) is constructed for parallel GICA+ICA by calculating the L_2_-distance between group- and subject-specific spatial component maps, capturing subject component variability from a group template. This variability matrix is then utilized to allow multimodal associations to directly influence the GICA and ICA estimation iteratively, hence leveraging the full temporal information of the fMRI as well as spatial variance (**Fig. 1**). Another advantage of parallel GICA+ICA compared with current existing multimodal fusion methods is that it also enables post functional network connectivity (FNC) analysis for fMRI after identification of sMRI links from the fusion analysis as well as other post analysis such as spectra and dynamic information (Hutchison, et al., 2013) for fMRI.

**Figure. 1.**
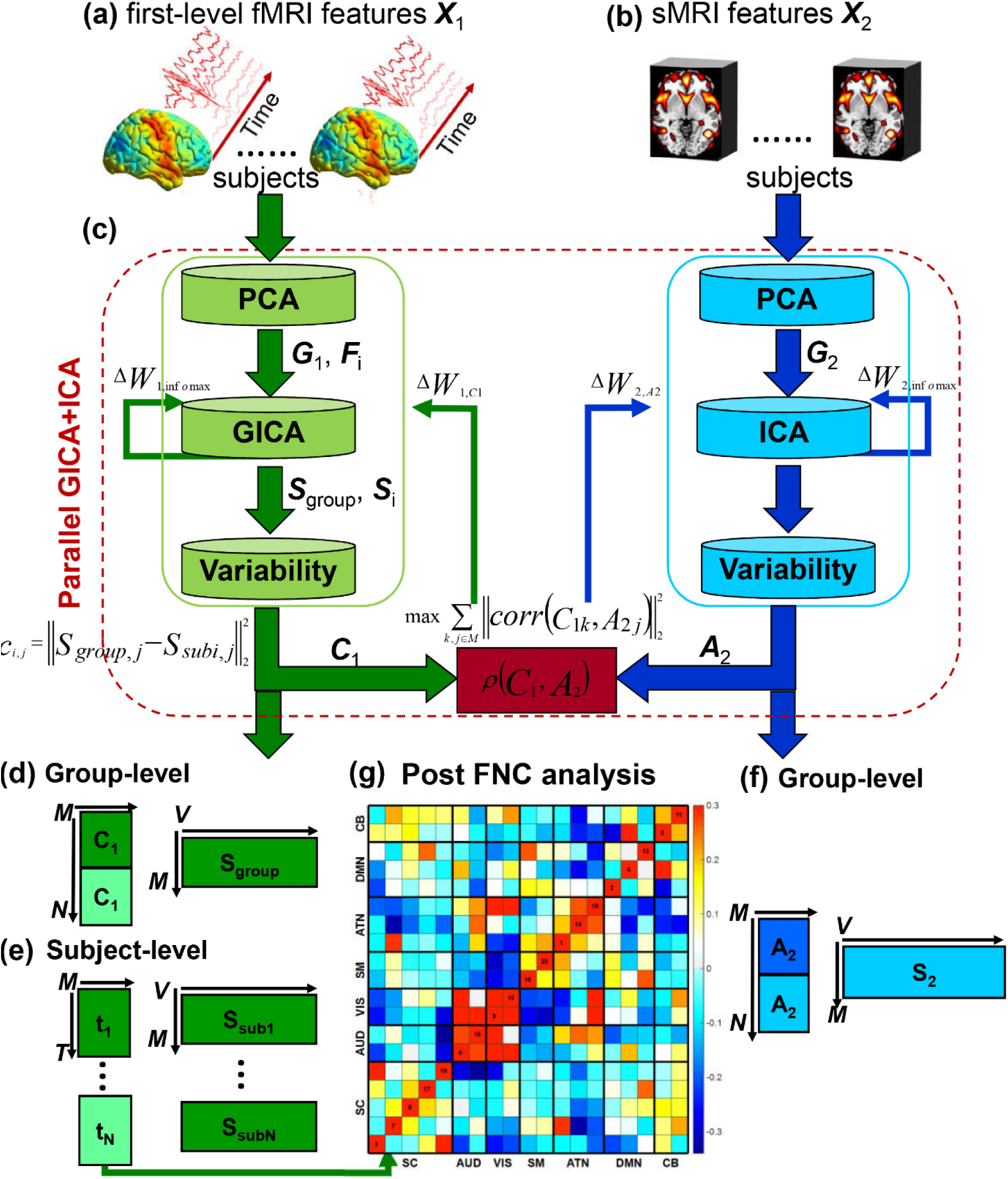
Flowchart of the proposed parallel GICA+ICA approach. (a) First-level fMRI features (***X***_1_) from preprocessed fMRI data (e.g. preprocessed spatiotemporal fMRI data). (b) SMRI features (***X***_2_) from preprocessed sMRI data (e.g. voxelwise gray matter volume or concentration). (c) Parallel GICA+ICA, which includes maximizing the independence for both modalities based on GICA and ICA portions separately, as well as maximizing the correlation between the variability matrix from GICA of fMRI and subject expression profiles from ICA of sMRI. (d) Group-level components and variability matrix obtained from GICA portion. (e) Subject-level components and time courses obtained from GICA portion. (f) Group-level components and subject expression profiles resulting from ICA portion. (g) Post functional network connectivity (FNC) analysis for fMRI time courses. PCA: principal component analysis; GICA: group independent component analysis; ***F***_*i*_: dewhitening matrix from subject-level PCA for fMRI; ***G***_1_: dewhitening matrix from group-level PCA for fMRI; ***G***_2_: dewhitening matrix from group-level PCA for sMRI; ***S***_group,j_: group components from GICA; ***S***_subi,j_: subject-specific components from GICA; ***C***_1_: variability matrix calculated by the L_2_-distance between group- and subject-specific GICA spatial maps; ***A***_2_: mixing matrix from sMRI, which also represents between-subject variability. Δ***W***_1,Infomax_ and Δ***W***_2,Infomax_ are the gradient updates obtained from GICA and ICA portions, separately. Δ***W***_1,C1_ and Δ***W***_2,A2_ are the gradient updates obtained from the between-modality linkage-regularized optimization. Dark and light colors in the variability matrix in panels (d-f) represent two groups, for example, schizophrenia patients and healthy controls.

In psychopathological studies, mounting evidence shows that schizophrenia (SZ) is associated with abnormal FNC between different brain networks, for example, visual, auditory and default mode networks (Friedman, et al., 2008). Based on the proposed method, we can perform post FNC analysis to identify abnormal connections between brain networks that are also associated with structural covariation. In this paper, we aim to apply the proposed method to identify linked functional network variability and structural covariations, and ultimately predict cognitive scores based on the identified linked fMRI-sMRI features. The Function Biomedical Informatics Research Network (fBIRN) phase III datasets (n=311) (Keator, et al., 2016) were used as a discovery cohort and the Center for Biomedical Research Excellence dataset (Jorge Nocedal, 1999) (COBRE, n=177) were used as a replication cohort. Results show that the proposed method can extract linked fMRI-sMRI components pair in both simulation and real human brain data. Moreover, the identified linked fMRI-sMRI features can predict multiple cognitive scores of fBIRN cohort, which demonstrates the biomarker property of the multimodal features extracted by the proposed method. Furthermore, these identified linked fMRI-sMRI features can also predict multiple cognitive scores of an independent COBRE cohort, demonstrating the ability of parallel GICA+ICA to identify potential biomarkers and the wide utility of the proposed method for the study of brain disorders.

## 2. METHODS AND MATERIALS

The main idea of parallel GICA+ICA is straightforward. As shown in **Fig. 1**, in order to retain the temporal nature of fMRI, we define a new variability matrix (***C***_1_) to capture functional subject-wise variability by calculating the L_2_-distance between group-level ICs (***S***_*group,j*_) and subject-level ICs (***S***_*subi,j*_) (*i* = 1,2,…, N, *j* = 1,2,…, M, where N is subject number and M is component number), in correspondence to the subject expression profiles (***A***_2_) from sMRI. In addition to maximizing the independence of each component for each modality, a regularization term is added to simultaneously maximize the correlation between functional (***C***_1_) and structural (***A***_2_) between-subject variabilities, as shown in **Fig. 1(c)**.

### 2.1 Parallel group ICA + ICA

Assume that ***X***_1_ = [***x***_1_; ***x***_2_;…; ***x***_*N*_] is fMRI data that is concatenated over subjects in the temporal dimension, where ***X***_1_ is in dimension of NT × V, in which T is time point number, V represents voxel number, and ***x***_*i*_ is the T × V data matrix. First, principal component analysis (PCA) is performed on fMRI data (***X***_1_) for dimension reduction at subject- and group-levels respectively. Let 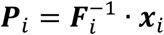 be the L × V dimension reduced data matrix of subject *i*, where 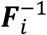 is the L × T subject-level whitening matrix (determined by subject-level PCA decomposition) and L is the rank of the PCA decomposition. The temporal dimension reduced data are then concatenated over subjects in the temporal direction and a group-level PCA is performed to further reduce the temporal dimension of the group data to the number of ICs M, as summarized in equation (1).

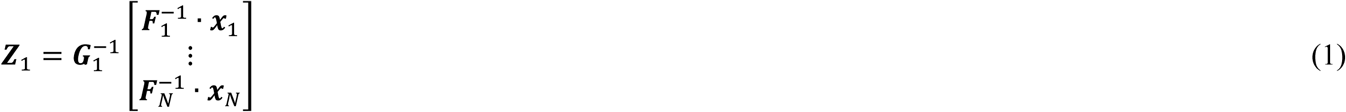

where 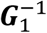 is the M × (N · L) group-level whitening matrix (determined by group-level PCA), ***Z***_1_ is the reduced data matrix for the fMRI modality.

After ICA decomposition, we can write ***Z***_1_ = ***A***_1_ · ***S***_*group*_, where ***A***_1_ is the M × M mixing matrix for fMRI and ***S***_*group*_ is the M × V group-level ICs. Substituting this equation for ***Z***_1_ into (1) and multiplying both sides by the group-level dewhitening matrix ***G***_1_, we have

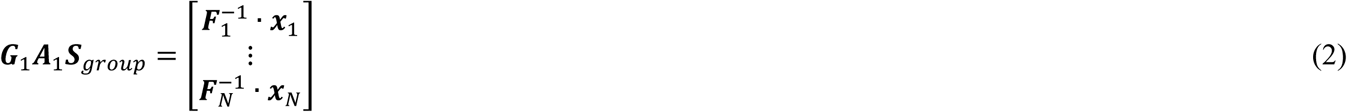

The matrix ***G***_1_ is partitioned by subject which provides the following expression

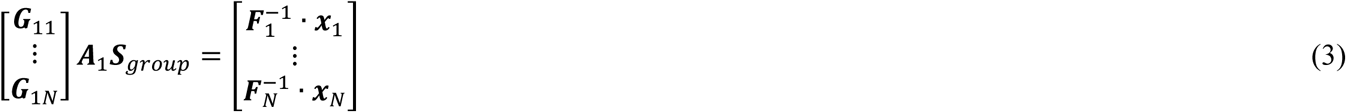

The equation for subject *i* by dealing only with the elements in partition*i* is as in (4)

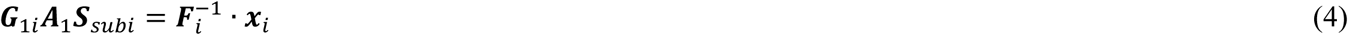

***S***_*subi*_ includes spatial maps for subject *i*. We now multiplying both sides of (4) by the subject-level dewhitening matrix ***F***_*i*_

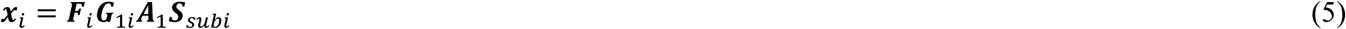

which provides the ICA decomposition of ***x***_*i*_ from subject *i*. ***S***_*subi*_ (M × V) contains M ICs and ***F***_*i*_***G***_1*i*_***A***_1_ (N × M) is subject-specific mixing matrix which contains the corresponding time course. This is the classic back-reconstruction formula from GICA (Calhoun, et al., 2001).

The between-subject functional variability matrix ***C***_1_ is defined as in (6) below:

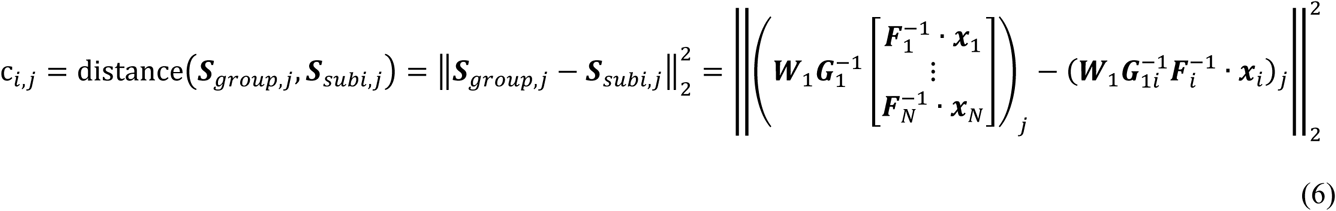

where ***W***_1_ equals the inverse of mixing matrix ***A***_1_.

Meanwhile, suppose that ***X***_2_ represents the sMRI data matrix, following the blind source separation theory, we can get (7).

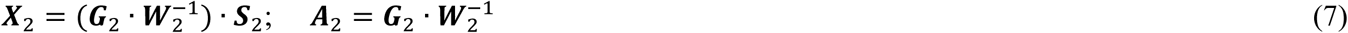

where ***G***_2_ is group-level dewhitening matrix for sMRI.

Thus we can get the final cost function for the proposed method, parallel GICA+ICA, as in (8):

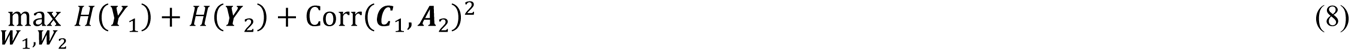

where

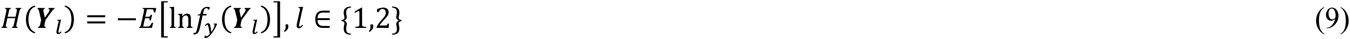

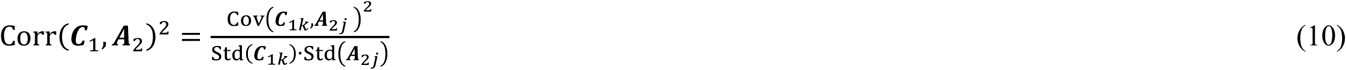

And

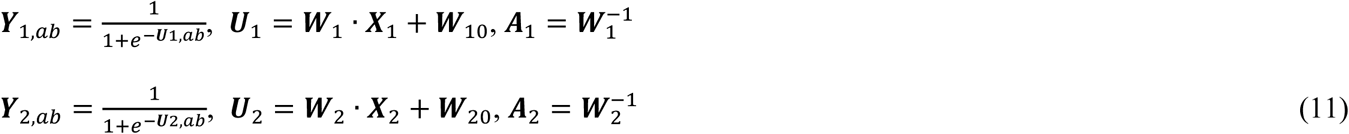

where ***Y***_*i,ab*_ and ***U***_*i,ab*_ represent elements for matrix ***Y***_*i*_ and ***U***_*i*_, respectively, with row index |*a*| and column index |*b*|. *H* is the entropy and Corr is the correlation. *k* and *j* represent the selected constrained ICs in each maximization iteration. Although the cost function (8) looks the same as pICA, we redefined a new variability matrix (6) here for fMRI. This matrix estimates the degree to which the subject specific component deviates from the cohort-common pattern, thus leads us to investigate whether this deviation may be associated with structural features. The first two terms in (8) are solved in parallel using the infomax framework (Amari, 1998). The third term is optimized using the steepest descent method, and the step size is calculated at each iteration on the selected ICs. Finally, we obtain the update rules as following:

For the first term (major updates for ***W***_1_):

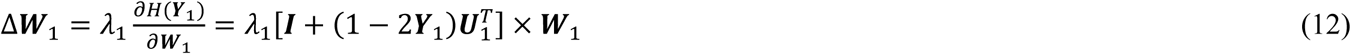

For the second term (major updates for ***W***_2_):

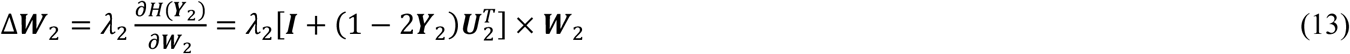

For the third term (minor updates for ***W***_1*k*_ and ***A***_2*j*_):

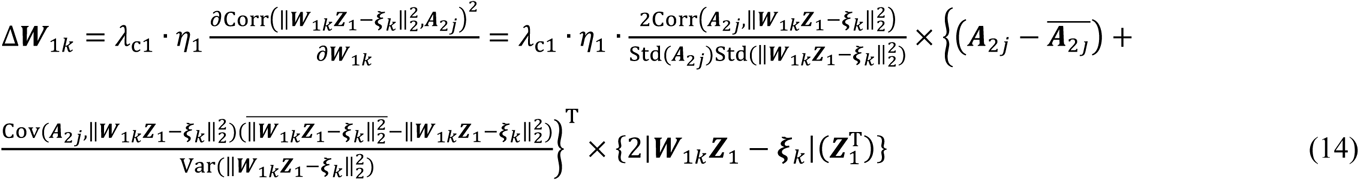

where 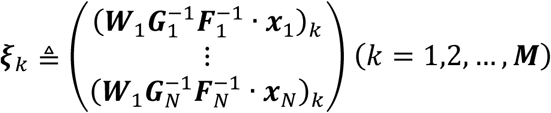

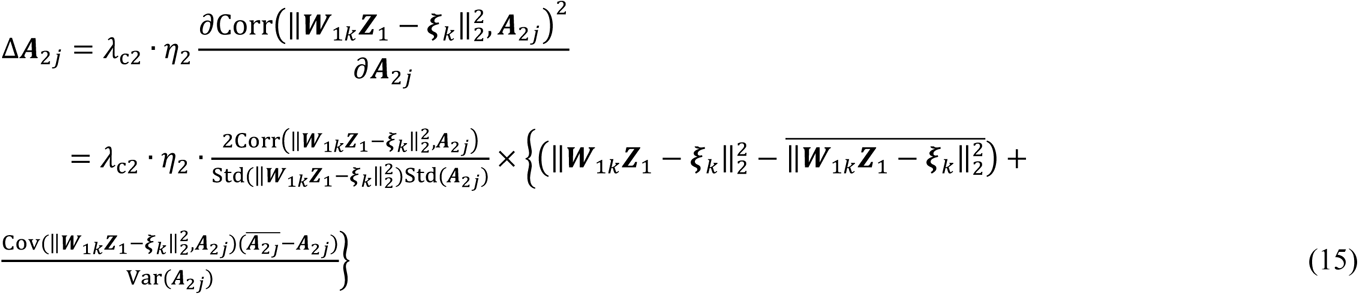

where *λ*_c*l*_ is the learning rate for fMRI, sMRI, and *η*_*l*_ is the step size calculated at each step according to Wolfe conditions (Jorge Nocedal, 1999). The learning rate plays an important role in algorithm convergence and balance between the modalities, as well as determines the weights during the maximization process.

The code of parallel GICA+ICA will soon be added in Fusion ICA Toolbox (FIT, http://mialab.mrn.org/software/fit).

### 2.2 Simulation

Then we simulated fMRI and sMRI data to compare parallel GICA+ICA with separate GICA and ICA to access the capability to detect accurate inter-modality linkage. Eight source maps were simulated using the simTB (Erhardt, et al., 2012) for each modality, which can be freely downloaded at http://mialab.mrn.org/software. The simulated brain networks were used as true sources ***S***_1*i*_ (in dimension of 100 × 100) for fMRI, each has 100 time points, and ***S***_2_ (in dimension of 150 × 150) for sMRI. TC matrix generated from simTB for fMRI is 100 × 8, mixing matrix ***A***_2_ for sMRI was randomly constructed in a size of 100 × 8. Variability matrix ***C***_1_ for fMRI is constructed by calculating the L_2_ norm between the group map and subject map, in which one column (the 1^st^ and 6^th^ column for ***C***_1_ and ***A***_2_) is carefully designed to be moderately (*r*=0.28) or highly correlated (*r*=0.49). Therefore, a linear mixture of ***TC***_*i*_ · ***S***_1*i*_ (***A***_2_ · ***S***_2_) will generate fMRI and sMRI data matrices of 100 samples with 10000 and 22500 voxels respectively. The observation matrix ***X***_2_ for sMRI is generated according to ***X***_2_ = ***I***_2_ + ***N***_2_ = ***A***_2_***S***_2_ + ***N***_2_ in which ***N***_2_ is the added noise which contains 15 peak signal-to-noise ratios (PSNR) noise levels. Here, the PSNR level is calculated from (16), which ranges from −10 dB to 17 dB. For fMRI, each ***x***_*i*_ is generated by ***x***_*i*_ = ***I***_*i*_ + ***N***_*i*_ = ***TC***_*i*_***S***_1*i*_ + ***N***_*i*_ (*i* = 1,2,…N). Here, for each specific PSNR, we randomly generated 10 same PSNR ***X***_2_ and ***x***_*i*_. Thus we can obtain the mean as well as the standard deviation of the inter-modality linkage estimation.

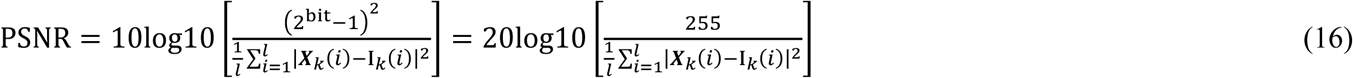

k = 1,2, bit = 8, *l* = 10000(for fMRI), 22500(for sMRI).

### 2.3 Real human brain data

For real human brain data, we used subjects collected from the fBIRN (Keator, et al., 2016) phase III study, including 149 SZ patients (37.9 ± 11.5) and 162 age-gender matched HCs (37.0 ± 11.0). All participants are adults between 18 and 65 years old. The Structured Clinical Interview for DSM-IV (SCID) (M. B. First, 2002) was used to diagnose patients by doctors. Furthermore, current or past psychiatric disorder or having a first-degree relative with an Axis-I psychotic illness HCs were excluded. The cognitive performance for both SZ and HC were measured by the Computerized Multiphasic Interactive Neurocognitive System (CMINDS) (van Erp, et al., 2015). There is no significant age (*p*=0.49) and gender (*p*=0.23) difference between SZ and HC for the fBIRN cohort.

The COBRE cohort consists of 94 SZs (35.6 ± 13.1) and 83 age-gender matched HCs (36.3 ± 12.5) and were assessed using a similar cognitive assessment battery, the Measurement and Treatment Research to Improve Cognition in Schizophrenia Consensus Cognitive Battery (MCCB) (Green, et al., 2004). Detailed cognitive information of fBIRN and COBRE subjects are presented in **Supplementary Table I-II**. There is also no significant age (*p*=0.45) and gender (*p*=0.91) difference for the COBRE cohort. Resting state functional MRI and structural MRI were obtained for both cohorts. Detailed imaging parameters and preprocessing steps can be found in **Supplementary “imaging parameters and preprocessing”** section. Written informed consent was obtained from all participants under protocols approved by the Institutional Review Boards for both fBIRN and COBRE cohorts.

## 3. RESULTS

### 3.1 Simulation results

The proposed parallel GICA+ICA algorithm was compared with separate GICA and ICA on carefully designed simulated data. One important property is that whether the proposed method can detect the inter-modality linkage of the target components accurately and significantly under both strong (*r*=0.49) and weak (*r*=0.28) inter-modality associations. **Fig. 2(a-b)** show the ability for estimating cross modality associations and corresponding significance for different fusion methods under 15 different noise levels with strong inter-modality linkage. The green line in **Fig. 2(b)** and **Fig. 2(d)** represents a significance level of *p*=0.05. It is clear that the proposed method, parallel GICA+ICA (red), could detect more accurate inter-modality linkage compared with separate GICA and ICA under strong real linkage. The estimation linkage accuracy trend goes down with high noise levels, which is also the same as in significance detection. **Fig. 2(c-d)** show the boxplot of estimating inter-modality associations and corresponding significance for different fusion methods under weak linkage. It is clear that parallel GICA+ICA outperforms separate GICA and ICA for weak linkage detection. More importantly, the association estimation of separate GICA and ICA decreases remarkably when the real correlation is weak (*r*=0.28), when compared with **Fig. 2 (a)** and **Fig. 2 (c)**, demonstrating the advantage of the proposed method in estimating association with weak linkage, the likely situation in real data. So collectively, the above results show that the proposed method provide more accurate detection of the inter-modality associations for both strong and weak links, and importantly, the fusion approach does not appear to inflate the multimodal links artificially.

**Figure. 2.**
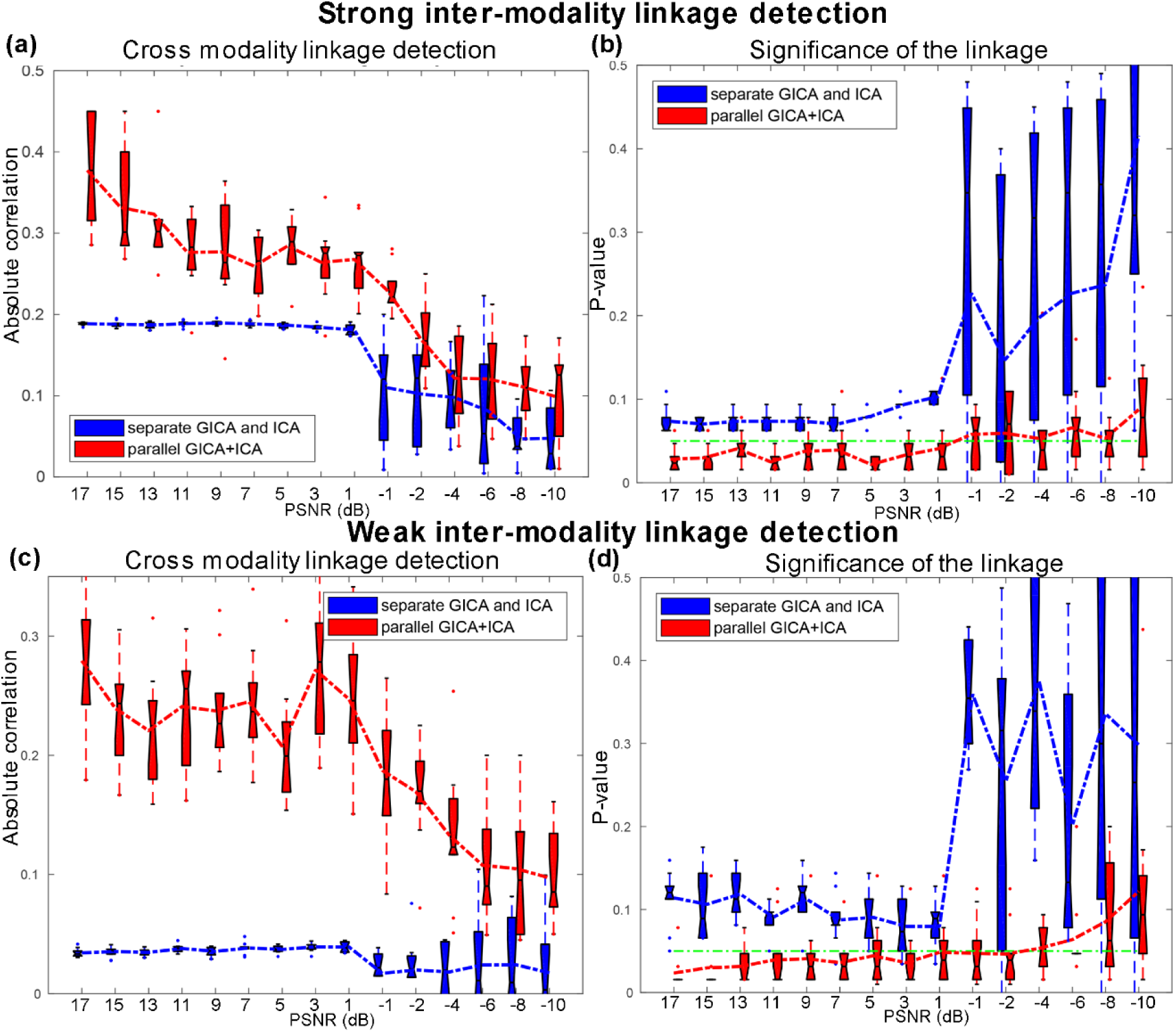
Comparison of parallel GICA+ICA (red) with separate GICA and ICA (blue) in a simulated two-way data fusion. (a-b) Cross modality linkage detection under strong association (*r*=0.49) and the corresponding significance *p* value. (c-d) Cross modality linkage detection under weak association (*r*=0.28) and the corresponding significance *p* value. The green lines in (b) and (d) represent *p*=0.05. These results show that, compared with separate GICA and ICA without fusion, parallel GICA+ICA yields more accurate (and significant) inter-modality associations regardless of whether the association is strong or weak, *i.e.* it improves the estimation of existing links while not inflating the link artificially.

Besides the linkage estimation, we also compared the source and mixing matrix (or variability matrix) estimation, as shown in **Fig. 3**. The estimation accuracy is defined as the correlation between the true mixing matrices/source(s) and the estimated mixing matrices/component(s). It is clear that parallel GICA+ICA can achieve high estimation accuracy of subject variability within the fMRI modality (**Fig. 3a**), and comparable estimation accuracy for structural mixing matrix and source, and also the subject specific functional source. Finally, the estimation accuracy for different IC numbers to extract modality linkage (under strong linkage, *r*=0.49) between target ICs is presented in **Fig. 4**. It is evident that parallel GICA+ICA out performs separate GICA and ICA in inter-modality linkage estimation.

**Figure. 3.**
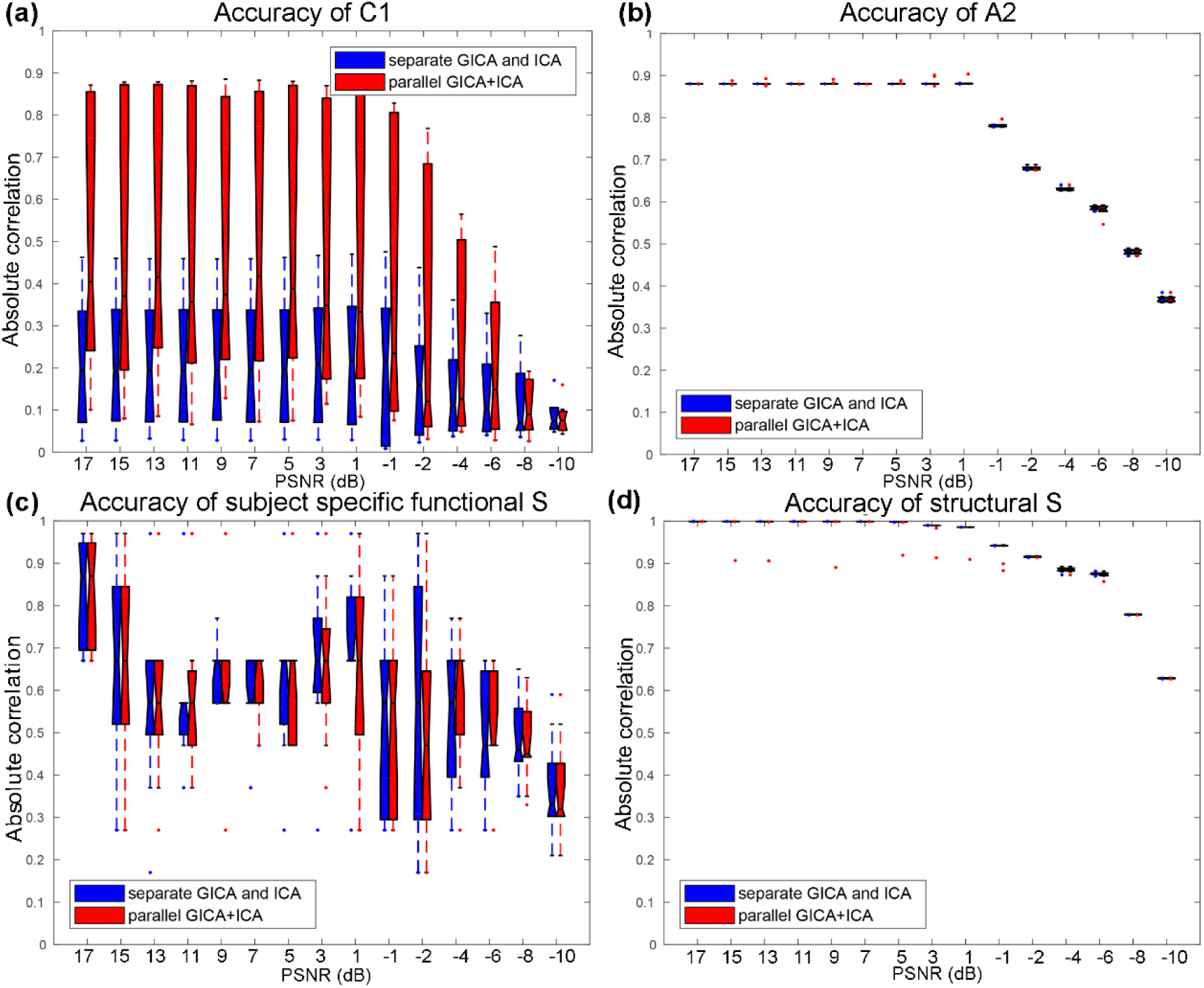
Estimation accuracy comparison for all the ICs under 15 level noises between parallel GICA+ICA (red) and separate GICA and ICA (blue). Results show that parallel GICA+ICA can achieve comparable estimation accuracy for both source and variability matrix.

**Figure. 4.**
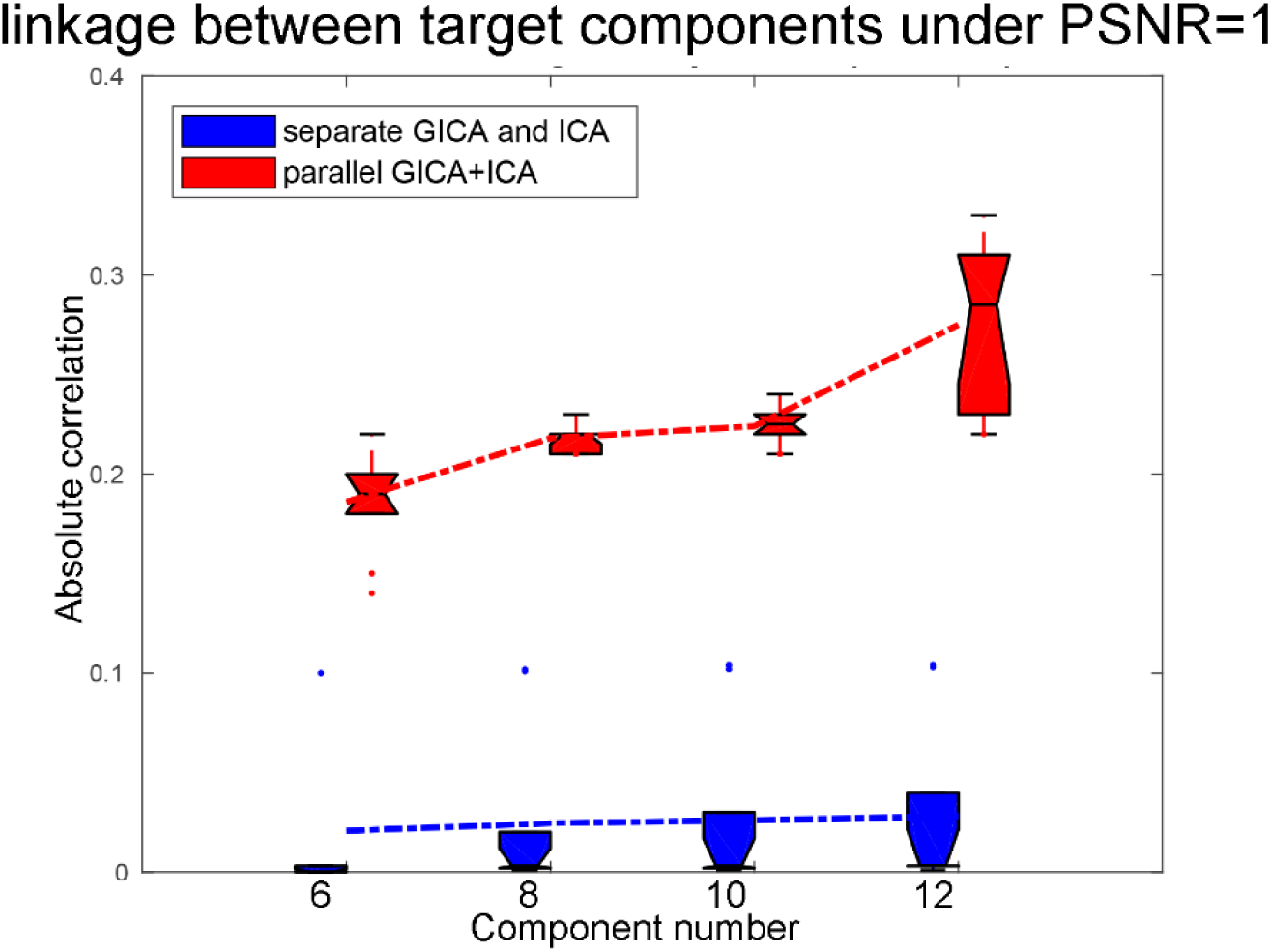
Comparison of inter modality linkage estimation when using different IC number *M*. The simulated true IC number = 8. Here we tested *M* that varies from 6 to 12. Parallel GICA+ICA outperforms separate GICA and ICA with regard to the inter modality linkage detection.

When determining the value of *λ*_c*i*_ and *η*_*i*_ in Eq (9) that control the weight between the independence constraint and the association constraint, we tune them adaptively. The criteria is that when the maximum value in ***W***_*i*_ is bigger than the predefined weight maximum (1.0 × 10^8^), then *λ*_c*i*_(*η*_*i*_) was update as *λ*_c*i*_(*η*_*i*_) = 0.95 · *λ*_c*i*_(*η*_*i*_) to avoid the blown up of ***W***_*i*_. This means that keeping the independence among components is always the first aim, the second is to maximize the cross modality correlation simultaneously.

### 3.2 Results of real human brain data

#### 3.2.1 Linked components pair

We also applied the proposed approach on fBIRN multimodal datasets including 311 subjects (162 HCs, 149 SZs). The original preprocessed TCs×voxels from resting-state fMRI, GM volume from sMRI were extracted and combined by parallel GICA+ICA to identify associated fMRI-sMRI components. The subject-level component was 50, which needs to be higher than the group level (Erhardt, et al., 2011), and the group-level component was 20 based on MDL (Li, et al., 2007) criterion for fMRI modality. Then two-sample t-tests was performed on the mixing coefficients as well as the variability loadings of each IC for each modality to compare patients and controls.

Among the 20 derived ICs, the 3^th^ IC of fMRI, the 19^th^ and the 3^th^ IC of sMRI were found to be the linked components pair (*r*=0.24, *p*=2.6e-05* between fMRI_IC3 and sMRI_IC19). * means FDR corrected for multiple comparisons. Note that this association still exist even after regressing out diagnosis (*r*=0.21, *p*=0.0002*). The mixing coefficients (or variability loadings) of the paired components show significant group difference (*p*=7.0e-06*, *p*=0.01) for fMRI_IC3 and sMRI_IC19. **Fig. 5(a)** shows the Z-scored brain maps (visualized at |Z| *>* 2), **Fig. 5(b)** presents the group difference of variability loadings for each modality, as well as the correlation between variability loadings and cognitive scores. So the red brain regions in sMRI denote a higher weights in HCs than SZs and the blue brain areas are opposite. While for fMRI component, the red regions indicate a higher variability compared with fMRI_IC3 in HCs than SZs and the blue regions are opposite.

**Figure. 5.**
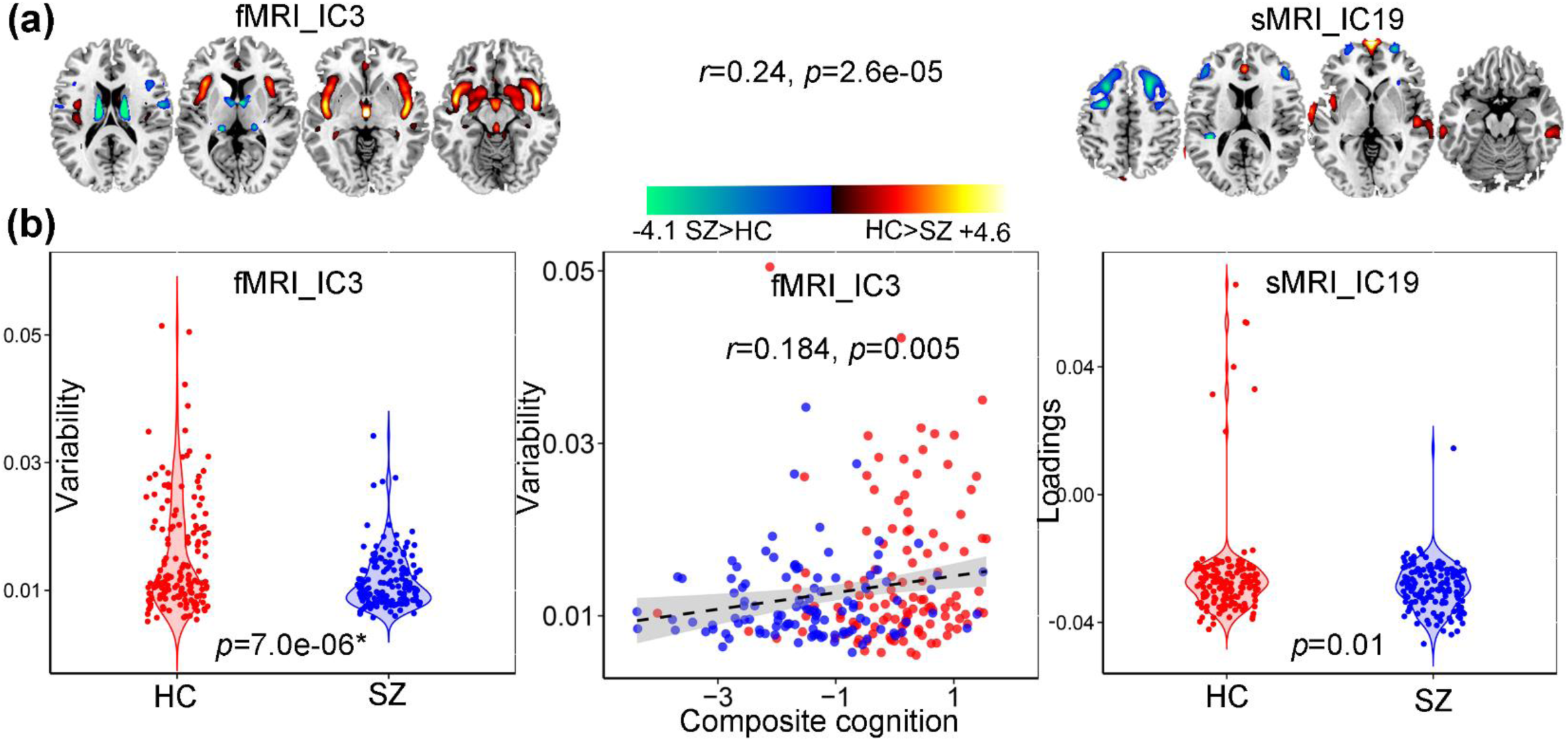
Linked components pair that indicate significant group differences in both fMRI and sMRI. (a) The brain maps of the identified linked component pair visualized at |Z|>2. (b) Group difference of the loadings for linked IC pair, and correlation between variability loadings of fMRI_IC3 and the CMINDS composite cognitive scores.

In addition, we found that fMRI_IC3 also correlated with sMRI_IC3 (*r*=0.14, *p*=0.01; *r*=0.12, *p*=0.051 after regressing out diagnosis). Significant group difference (*p*=0.001*) exist for the variability loadings of sMRI_IC3, which is also positively with speed of processing (*r*=0.349, *p*=4.3e-08*), working memory (*r*=0.344, *p*=6.1e-08*), visual learning (*r*=0.336, *p*=1.5e-07*), verbal learning (*r*=0.337, *p*=1.5e-07*) and composite cognitive scores (*r*=0.30, *p*=4.1e-06*), as displayed in **Fig. 6**. Note that after regressing out group label, the correlation between loadings of sMRI_IC3 and cognitive scores still remain significant. Details are presented in **Supplementary “regression out diagnosis”** section.

**Figure. 6.**
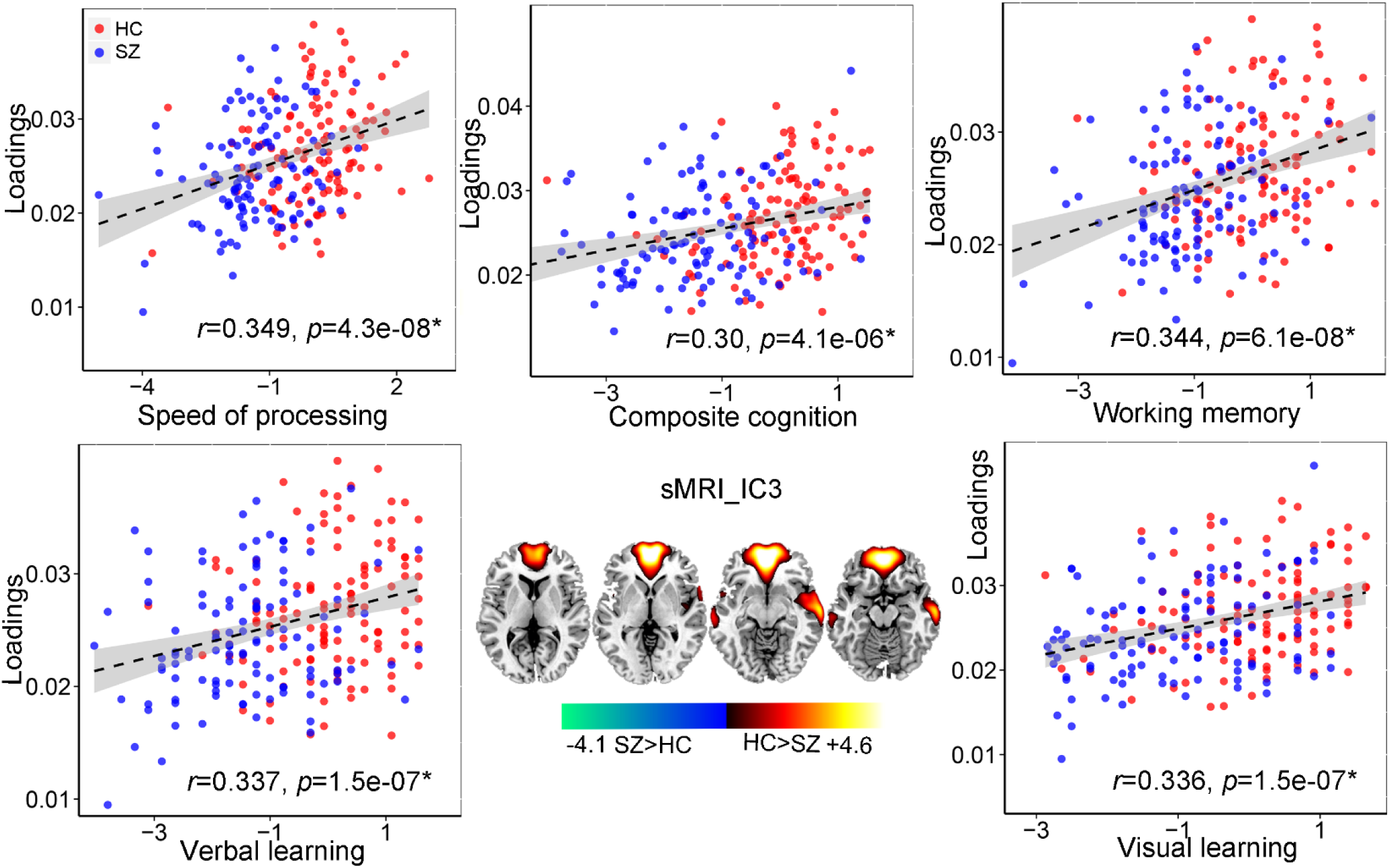
Correlation analysis between loadings of sMRI_IC3 and multiple CMINDS cognitive domains.

**Figure. 7.**
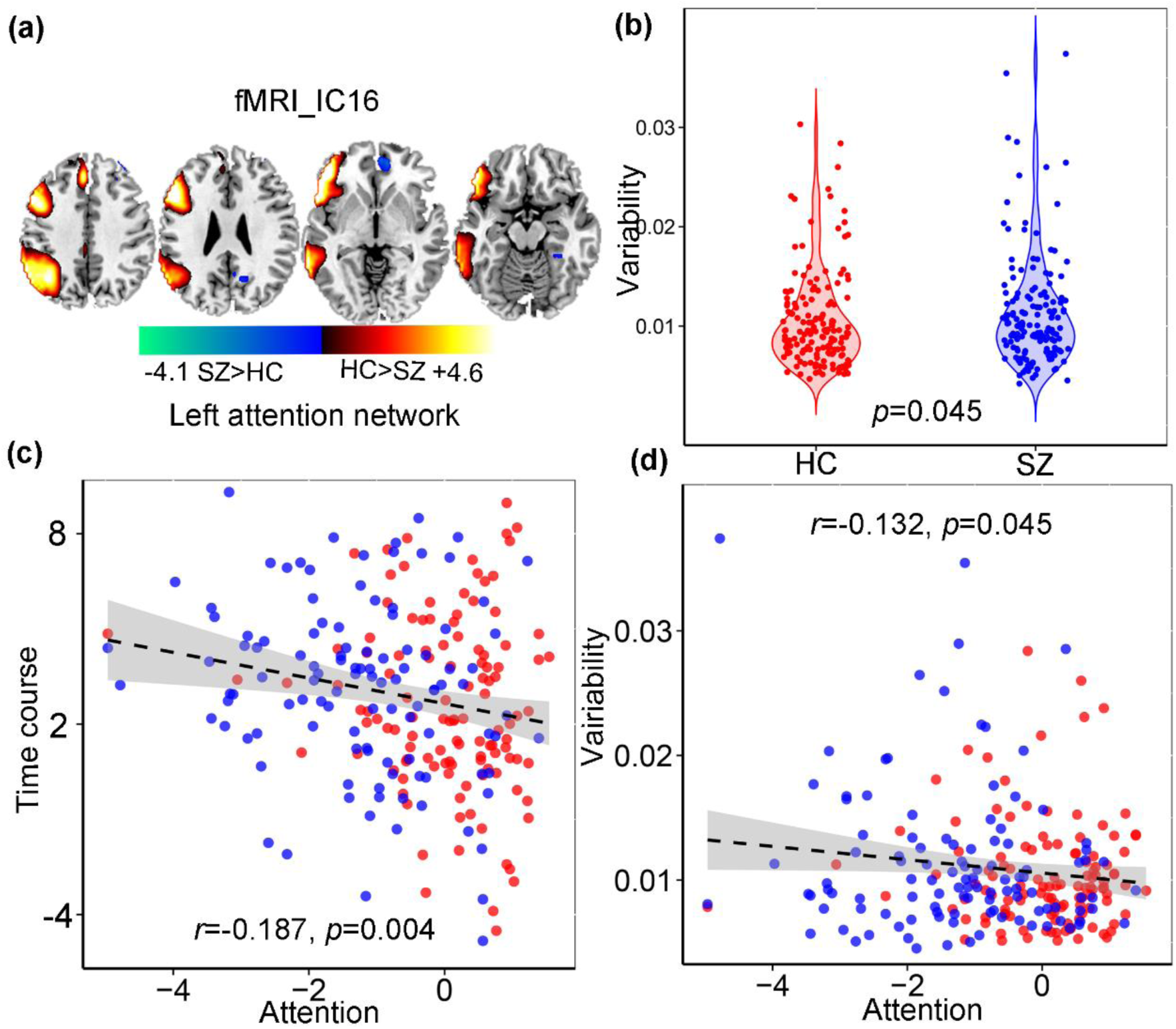
Modality specific and group discriminative left attention network in fMRI_IC16. (a) The brain maps visualized at |Z|>2. (b) Group difference of the loadings for fMRI_IC16. (c) Correlations between time courses of fMRI_IC16 and the CMINDS attention scores (HC: red dots, SZ: blue dots. (d) Correlations between variability loadings of fMRI_IC16 and the CMINDS attention scores.

#### 3.2.2 Modality specific group-discriminative IC

Apart from the linked components pair, we also identified one fMRI component IC16 containing the left attention network that shows marginal group difference (*p*=0.045), and the corresponding variability loadings is negatively correlated with attention subdomain (*r*=-0.132, *p*=0.045). More importantly, the time courses of fMRI_IC16 were also anti-correlated with attention scores (*r*=-0.187, *p*=0.004).

#### 3.2.3 Post FNC analysis

One of the advantages of the proposed method compared with other fusion methods is that after the fusion analysis, we can still perform FNC analysis using time courses of the corresponding components. Collectively, 19 components were selected from 20 components, with one artefactual component excluded. **Fig. 8(a-b)** shows the mean FNC matrix for HCs and SZs. The black lines partition the FNC matrix into the 7 categories: visual (VIS), default-mode network (DMN), auditory (AUD), attention network (ATN), sensorimotor (SM), cerebellar (CB) and sub-cortical (SC). We also compared the group difference of FNC matrix between HCs and SZs. Values in FNC matrix are calculated as −log10(*p*) × sign(*t*). Results show that the AUD-VIS, AUD-ATN, DMN-SC are group discriminating FNCs when *p* was thresholded as *p* < 0.01, while the AUD-VIS still remains significant group difference when *p* was thresholded as *p* < 0.0001 (FDR corrected).

**Figure. 8.**
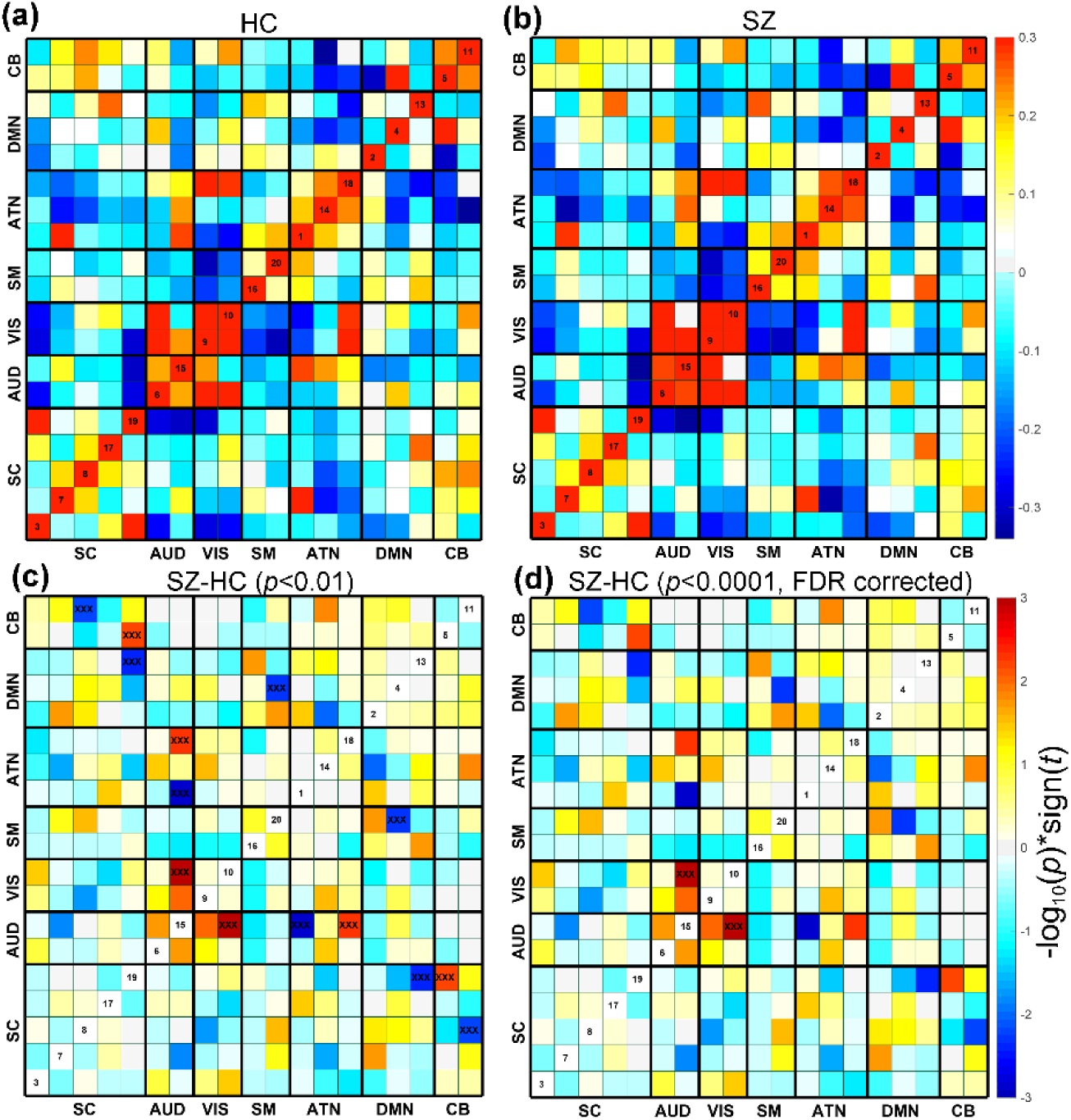
Post FNC analysis based on the fMRI results from parallel GICA+ICA. Mean FNC matrix for HCs (a) and SZs (b). (c-d) The group difference (SZ–HC) in FNC.

#### 3.2.4 Predicting ability

An ultimate goal of using imaging biomarkers is whether these identified imaging features is predictive on cognition or symptoms (Gabrieli, et al., 2015). To verify the predictability of the identified linked multimodal brain features by our proposed method, we used the extracted linked components ROIs (mean time courses in positive and negative brain networks in fMRI_IC3, and positive and negative brain networks in sMRI_IC19 and sMRI_IC3, details can be found in **Supplementary “Predictive biomarker extraction”** section) to predict multiple CMINDS cognitive scores. Based on the following Eq (17), a multiple linear regression for the cognitive scores was performed.

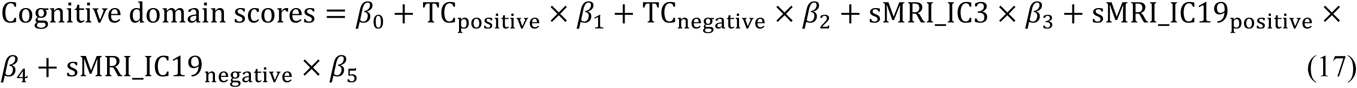

The predictive accuracy is measured by correlation between the estimated cognitive scores and its true values, as well as the normalized root mean squared prediction error (NRMSE) (https://en.wikipedia.org/wiki/Root-mean-square_deviation). Multiple cognitive domain prediction results are shown in **Fig. 9**. The five features identified based on the proposed parallel GICA+ICA were able to predict working memory (*r*=0.266, *p*=4.5e-05; NRMSE=0.19), verbal learning (*r*=0.254, *p*=8.0e-05; NRMSE=0.21) and composite cognitive scores (*r*=0.261, *p*=6.7e-05; NRMSE=0.15). More importantly, after regressing out diagnosis, the prediction still keeps significant for verbal learning (*r*=0.157, *p*=0.016) and composite cognitive domains (*r*=0.15, *p*=0.02), demonstrating the effectiveness of the proposed method in identifying linked multimodal biomarkers associated with brain disorders. To test the predictability of the extracted five linked multimodal features by the proposed method on new unseen dataset, we then extracted the brain features in COBRE cohort through fBIRN identified masks and predict an independent cohort(COBRE)’s multiple MCCB cognitive scores for working memory (*r*=0.235, *p*=0.001; NRMSE=0.15), verbal learning (*r*=0.25, *p*=8.5e-04; NRMSE=0.17), and composite cognitive scores (*r*=0.256, *p*=9.2e-04; NRMSE=0.18), as displayed in **Fig. 10**. Note that MCCB and CMINDS are similar but not identical cognition assessment batteries (van Erp, et al., 2015); hence, the cross-cohort prediction is a strong evidence to demonstrate the predictability of the identified multimodal features detected by parallel GICA+ICA. Furthermore, after regressing out diagnosis, the prediction still keeps significant, details are presented in **Supplementary “regression out diagnosis”** section.

**Figure. 9.**
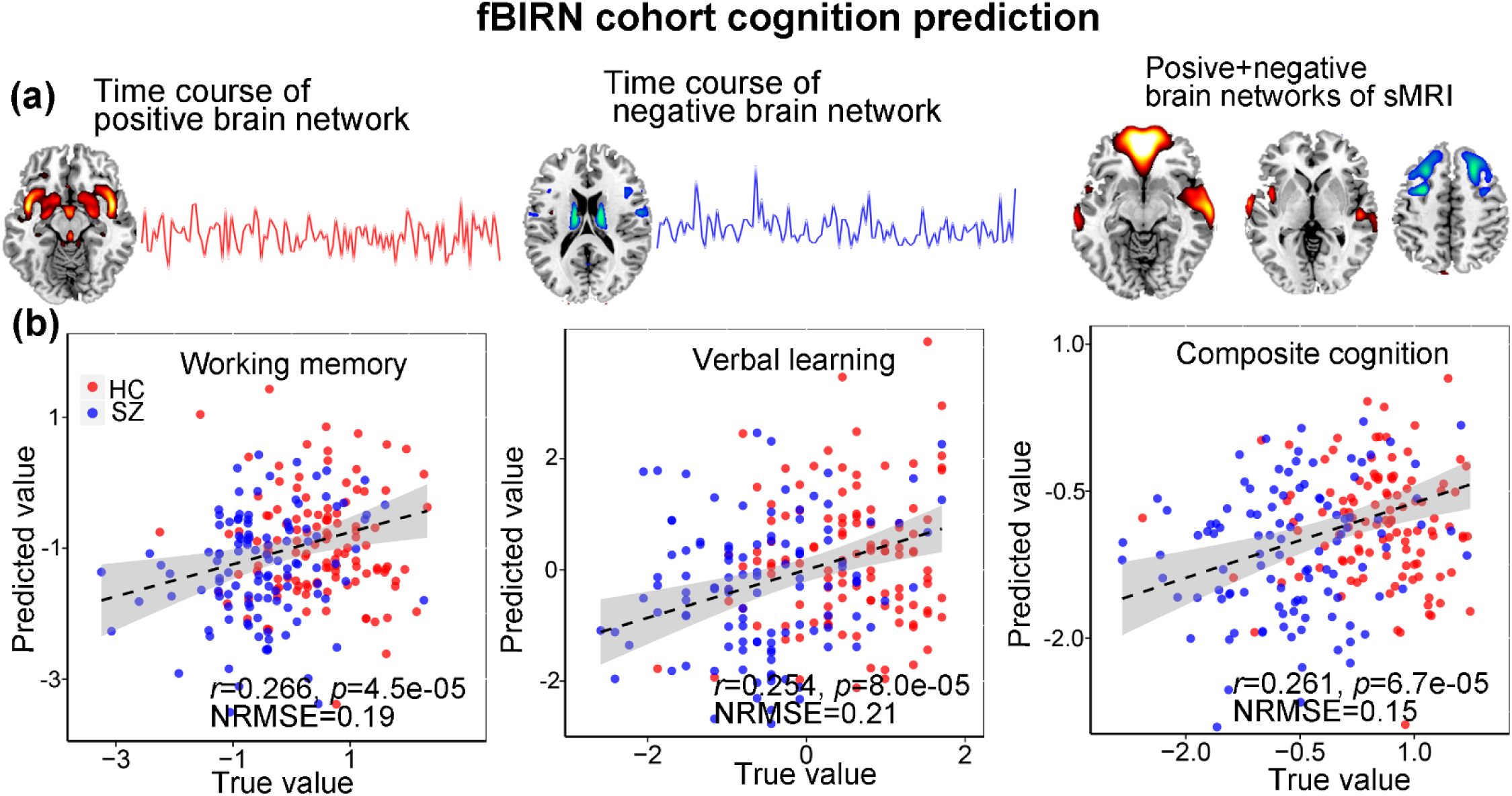
The identified linked fMRI-sMRI biomarkers and the prediction results on multiple cognitive and symptom scores of fBIRN cohort. Five modality-specific time courses and brain networks (a) were used as regressors in the multiple linear regression models to predict cognitive scores (b).

**Figure. 10.**
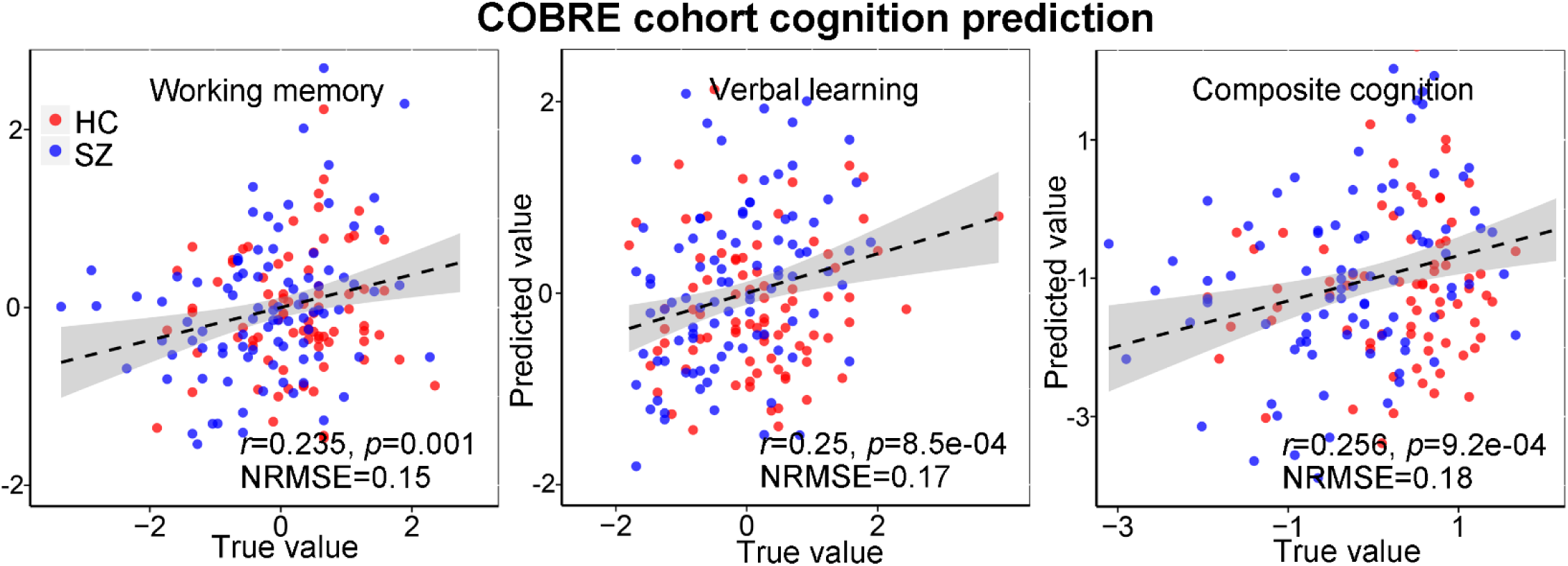
Prediction of multiple cognitive scores of COBRE cohort based on the identified multimodal features in the linked components pair by pGICA+ICA in fBIRN cohort.

## 4. DISCUSSION AND CONCLUSION

In this paper, we proposed a novel temporal information included parallel GICA+ICA, by adding a constraint that maximize the correlation between two variability matrices from fMRI and sMRI, aiming to extract the associated functional network variability with sMRI covariations. Compared with traditional multimodal fusion methods (mCCA, jICA, pICA, pICAR, mCCA+jICA, MCCAR+jICA, linked ICA), parallel GICA+ICA can deal with first-level features (temporal information included) in fMRI related fusion analysis. Another advantage is that we can also perform FNC analysis based on the fMRI results of parallel GICA+ICA. To the best of our knowledge, this is the first established method to incorporate temporal information in multimodal fusion analysis, which provides a new perspective to interpret interrelated patterns between functional network variability and co-varied structural measures.

In simulations, we compared parallel GICA+ICA with separate GICA and ICA on the performance of detecting linked fMRI-sMRI components. Our results indicate that parallel GICA+ICA provides more accurate detection of inter-modality linkage regardless of whether it’s strong or weak, which means that it doesn’t inflate the link artificially as well as achieves comparable accuracy on both source maps and mixing coefficients. More importantly, the association estimation of separate GICA and ICA decreases remarkably when the real correlation is weak (*r*=0.28) compared with parallel GICA+ICA, as shown in **Fig. 2(c-d)**, demonstrating the advantages of the proposed method in estimating association with weak linkage, which is always the case in real data.

In the real multimodal brain imaging fusion application, we combined data from gray matter volume and brain function from SZ patients and HCs. One linked component pair (fMRI_IC3-sMRI_IC19) was identified that show significant group difference between SZ and HC. Subcortical regions including bilateral insular, striatum, thalamus and hippocampus are identified in the linked fMRI_IC3. These regions have been widely reported as dysfunctional brain areas in SZ (Friedman, et al., 2008), and also associated cognitive deficits in SZ (Bush, et al., 2005; Glahn, et al., 2005). SMRI_IC3 is correlated with several cognitive domains, which is consistent with the prefrontal cortex detected in sMRI in our results being involved in multiple high-order cognitive functions including manipulating and maintaining information in problem solving (Minzenberg, et al., 2009), working memory (Aleman, 1999; Potkin, et al., 2009) and decision making (Barch and Ceaser, 2012). We also identified one modality-specific fMRI component containing the left attention network, whose variability loadings show group difference and also correlated with attention domain. More importantly, the corresponding time courses of fMRI_IC16 are also correlated with attention domain. The identified correspondence between the identified brain areas and the correlation with attention demonstrates well the effectiveness of the proposed method. Apart from the traditional multimodal fusion analysis, we also performed FNC analysis, and identified one abnormal FNC pair between visual and auditory networks compared with SZ and HC, which is widely reported associated with the auditory hallucination impairment with SZ (Friedman, et al., 2008; Liu, et al., 2018).

Finally, the primary goal of brain imaging studies is to identify biomarkers that can predict individual cognition or symptoms scores (Gupta, et al., 2015). Based on the identified multimodal brain features in the linked component pair, multiple cognitive scores can be predicted in fBRIN cohort. These features can also be applied to independent cohort (COBRE) to predict unseen subjects on similar but different cognitive measures. All the above results demonstrate the ability of parallel GICA+ICA in detecting associated multimodal components pair that contains potential biomarkers related with mental disorders, suggesting a wide utility in the brain imaging community (Carter, et al., 2008; Jiang, et al., 2018).

In addition to sMRI, other modalities could also be fused with temporal information included fMRI based on the proposed method, such as fractional anisotropy from dMRI (Sui, et al., 2018). Parallel GICA+ICA can be applied straightforwardly to study other brain disorders, such as bipolar disorder, depression (Etkin and Wager, 2007; Qi, et al., 2018b), and attention-deficit/hyperactivity disorder, suggesting a comprehensive application in brain disorder related imaging studies. Moreover, except for the distance that measuring the subject variability, other measures (such as correlation or covariance between group and subjects) can be directly incorporated into our proposed method as well. Furthermore, apart from static FNC analysis in the current study, dynamic FNC (Calhoun, et al., 2014) can also be calculated based on the parallel GICA+ICA results.

In summary, this study proposed a novel temporal information added multimodal fusion method, *i.e.*, parallel GICA+ICA, and verified its effectiveness in both simulation and real human brain imaging data. To the best of our knowledge, this is the first proposed method that can directly link first-level fMRI features with sMRI data, seeking for potential linked multimodal MRI markers in brain disorders. Based on the proposed parallel GICA+ICA, we identified one linked fMRI-sMRI pair that was indicated to be associated with major SZ deficits in multiple reports, which can also be used to predict multiple cognitive scores of fBIRN cohort, as well as an independent COBRE cohort, promising a wide usage of the proposed method in detecting potential linked multimodal biomarkers for psychosis.

## ACKNOWLEDGEMENTS

This work was supported by the NIH grants R56MH117107, R01EB005846, 1R01MH094524, P20GM103472, and P30GM122734 as well as NSF grant 1539067, and the Natural Science Foundation of China No. 81471367, 61773380, the Strategic Priority Research Program of the Chinese Academy of Sciences (CAS) (grant No. XDB03040100).

